# Radiotherapy Technique Determines the Magnitude and Persistence of Lymphocyte DNA damage

**DOI:** 10.1101/2025.05.07.652586

**Authors:** Zsuzsa S. Kocsis, Péter Ágoston, Gyöngyi Farkas, Gábor Székely, Gyöngyvér Orsolya Sándor, Kliton Jorgo, László Gesztesi, Tibor Major, Csilla Pesznyák, András Herein, Gábor Stelczer, Dalma Mihály, Georgina Fröhlich, Zoltán Takácsi-Nagy, Csaba Polgár, Zsolt Jurányi

**Author notes:** **Corresponding author:** Dr. Zsolt Jurányi, 1122, Ráth György utca 7-9. Budapest, Hungary; /1379.

## Abstract

Data comparing the biological dose of radiotherapy techniques in the same patient group are scarce. Furthermore, the assessment of lymphocyte damage caused by radiotherapy can be of importance as immunotherapies are used more frequently.

By applying the chromosome aberration technique, over five years prospective comparison of the biological impact of four different types of treatments for low- and intermediate-risk prostate cancer was performed (at 3, 6, 9, 12, 24, 36, 48, and 60 months, 195 patients): conventional LINAC (linear accelerator) (70–78 Gy), CyberKnife (40–37.5 Gy) teletherapy, low-dose rate brachytherapy (LDR; 145 Gy) and high-dose rate brachytherapy (HDR; 19–21 Gy). Multivariate regression analyses were performed to analyze the predictive potential of chromosome aberrations for side effects. The median follow-up was 60 months.

We found that teletherapy techniques (conventional LINAC and CyberKnife therapy) caused 1.7–3.2-fold more chromosomal aberrations than brachytherapy did. At three months, 4.6–12.7% of the lymphocytes were damaged. Five years after treatment, the total aberration values of conventional LINAC and LDR brachytherapy patients were still significantly greater than those before therapy (p=0.035 for LINAC and p=0.029 for LDR BT). We found that significant regression models suggest total aberrations or aberrant cell frequency might predict side effects in addition to biological effective dose (BED) and irradiated volume (V_100%_) (p=0.005 for model including total aberrations, p=0.003 for aberrant cell frequency).

We reported a lower biological dose and fewer side effects in brachytherapy patients. We also demonstrated that long-term lymphocyte damage was dependent on the type of radiotherapy.

**Novelty and impact:** We analyzed radiation dose of the immune cells after four types of radiotherapy for five years. The damage of the lymphocytes can be of significance, if patients receive immunotherapy after radiotherapy in the future. We demonstrated chromosome aberrations were still present in lymphocytes years after the radiation. We found that there can be threefold difference in lymphocyte damage between therapy types. We also showed capacity of chromosome aberration technique to predict side effects.

## Introduction

Approximately half of all patients with tumors are treated via different radiotherapy techniques ^1^. Treatment modalities and fractionation schedules are commonly evaluated using the linear– quadratic model to describe dose–response relationships and via biological effective dose calculations to compare different fractionation schemes. However, the relative contribution of for example radiation energy, fractionation, overall treatment time, dose rate to the overall effect on healthy cells remain uncertain, limiting deeper mechanistic understanding^2^.

Because radiation-induced cellular biological DNA damage is commonly quantified using lymphocytes as surrogate markers of healthy tissue ^3^, it also describes the condition of immune cells. DNA damage was shown to cause T-cell senescence ^4-6^ and senescence of lymphocytes was demonstrated to be a factor in the success rate of immunotherapy ^7-9^. Prostate cancer is one of the few tumor types, that can be routinely treated through multiple different radiotherapy techniques, therefore, we used it as a model for this dual purpose, even if immunotherapy is not a major treatment modality in this tumor site.

In our institute, teletherapy can be delivered by conventional linear accelerators (LINACs) or by a CyberKnife LINAC with the precision of the robotic arm of the radiotherapy machine. Higher conformality allows higher doses per fraction and shorter overall treatment times. On the other hand, brachytherapy employs radioactive isotopes implanted into the tumor. The movement of the tumor causes less difficulty in this case, and smaller safety margins around the clinical target volume (to account for organ motion) can be applied. A high dose gradient also helps prevent normal tissue toxicity. We compared the gold standard values of biological dosimetry, chromosomal aberration frequencies, following treatment with these four radiotherapy modalities.

The chromosomal aberrations (CAs) are also suggested to predict radiotherapeutic side effects as surrogate markers (without being causative) ^10-12^ for which there is no universally accepted standard method in clinical practice. We tested this hypothesis by comparing chromosomal aberrations and graded urogenital and gastrointestinal radiation toxicities.

Furthermore, there is also scarce information about how long it takes for damaged cells to be eliminated ^13^ and we found years-long persistence of chromosome aberrations.

## Patients and methods

### Patients

Between April 2015 and June 2021, low- and intermediate-risk prostate cancer patients ^14^ (N=196) were enrolled in our study. Patients with previous tumors were excluded. Blood samples were taken before therapy and at the follow-up visits, when PSA measurement was also performed. The physicians evaluated patients’ side effects according to the RTOG-EORTC grading system, and the International Prostate Syndrome Score (IPSS) and the quality of life (QoL) questionnaires were completed by the patients themselves. Blood samples were taken once from healthy men (N=22) whose age matched that of the patient group. We performed the study according to the Helsinki guidelines with the approval of the Ethics Committee (16738-2/2015/EKU). Both patients and volunteers signed informed consent forms.

### Radiotherapy

At our institute, four types of radiotherapy are available for nonmetastatic prostate cancer patients.

Brachytherapy techniques are suggested for patients whose prostate measures less than 60 cm^3^, whose rectum–prostate distance is greater than 5 mm, and who have no pubic arch interference. Patients with an initial IPSS greater than 15 were advised against brachytherapy, because one-fraction HDR monotherapy and LDR brachytherapy were given in a randomized clinical study (NCT02258087). Otherwise, patient preference for teletherapy or brachytherapy was considered. The dose ⍰ volume constraints are listed in Supplementary Table 1. The CyberKnife machine was installed at our institute in 2018. Before that, all teletherapy patients received LINAC therapy, after the installation patients could also opt for CyberKnife therapy.

LINAC therapy: For intermediate-risk patients, the prostate CTV was expanded with a 0.5 cm margin (except in the posterior direction, where it was not extended), and a 1.0 cm margin was added cranially for seminal vesicle irradiation. Low-risk patients received 70–78 Gy to the prostate, and intermediate-risk patients received 57.4–60 Gy to seminal vesicles. Both simultaneous integrated boost (SIB: 2.5 Gy/fraction on the prostate, 2.05 Gy/fraction on the seminal vesicle) and classical (2 Gy/fraction) fractionation were acceptable ^15^. Treatment planning was performed via Eclipse v13.7 (Varian, Palo Alto, USA) or Pinnacle v9.8 (Philips, Eindhoven, The Netherlands) systems. The dose-volume constraints are shown in Supplementary Table 1.

Cyberknife therapy: For a precise setup, four gold markers were inserted into the prostate two weeks prior to the planning CT. The prescribed dose was 37.5–40 Gy to the prostate, and for intermediate-risk patients, 30–32.5 Gy was also given to the seminal vesicles CTV. Radiotherapy was delivered via a Cyberknife machine (Accuray, Madison) in 5 fractions, which were administered every second workday ^16^.

LDR BT (seed BT): Low-dose rate brachytherapy was performed with stranded seeds (^125^I seeds (BEBIG, Germany)). After anesthesia, catheters were inserted into the prostate with rectal ultrasound guidance in the lithotomy position, and the seeds were implanted (SPOT PRO 3.1 and Oncentra Prostate 3.2.2). 145 Gy was delivered on the surface of the prostate. ^17,18^

HDR BT: After Foley catheter insertion, transrectal ultrasound-guided needle insertion was performed, and an intraoperative plan was made (Oncentra Prostate 3.2.2). The dose delivery was executed with an afterloading machine (Elekta Brachytherapy, Veenendaal, The Netherlands), and the prescribed dose was 19 or 21 Gy (^192^Ir) ^19^.

### Baseline characteristics

The distribution of Gleason score, risk ^14^ and androgen deprivation therapy use varied significantly between the groups, but no disparities were observed in the T status, age or iPSA level (Table 1).

**Table 1:** Baseline characteristics of the patients in our study.

|  | conventional<br>LINAC | Cyberknife | LDT BT | HDR BT | p value |
| --- | --- | --- | --- | --- | --- |
| **Patients** | 25 | 51 | 68 | 52 |  |
| **Age**<br>**(Range)** | 68.5 (53-79) | 67.4 (54-78) | 67.0 (50-79) | 65.1 (53-75) | 0.065 |
| **iPSA**<br>**(range)** | 11.2 (3.0-20.0) | 9.6 (2.8-18.2) | 9.6(4.4-20.0) | 8.9 (4.9-18.5) | 0.153 |
| **T status** |  |  |  |  |  |
| T1b | 1 (4,0%) | 1 (2.0%) | 0 | 0 | 0.152 |
| T1c | 6 (24,0%) | 6 (11.7%) | 17 (25.0%) | 13 (25.0%) |  |
| T2a | 7 (28%) | 21 (41,2%) | 23 (33.8%) | 16 (30.8%) |  |
| T2b | 3 (12%) | 15 (29.4%) | 10 (14.7%) | 7 (13.4%) |  |
| T2c | 7 (28%) | 7 (13.7%) | 18 (26.5%) | 16 (30.8%) |  |
| n.a. | 1 (4%) | 1 (2.0%) | 0 | 0 |  |
| **Gleason score** |  |  |  |  |  |
| ≤6 | 12 (48.0%) | 21 (41.2%) | 48 (70.6%) | 36 (69.2%) | 0.001 |
| 7 | 13 (52.0%) | 30 (58.8%) | 17 (25.0%) | 16 (30.8%) |  |
| n.a. | 0 | 0 | 3 (4.4%) | 0 |  |
| **Risk** |  |  |  |  |  |
| low/inter. | 4/21 | 9/42 | 18/50 | 22/30 | 0.019 |
| HT0/HT1 | 2/23 | 36/15 | 23/45 | 25/27 | <0.0001 |
| SIB/conv. | 17/8 | 51/0 | - | - |  |
| 19/21Gy | - | - | - | 19/33 |  |

Patients characteristics are shown according to the different treatment groups. (iPSA =initial PSA, LDR BT=low dose rate brachytherapy, HDR BT= high dose rate brachytherapy, HT1= androgen deprivation hormone therapy receiving, SIB= Simultaneous integrated boost

### Lymphocyte sample preparation and chromosome analysis

Blood samples were collected via venipuncture into Li-heparinized tubes. Lymphocytes were induced to proliferate with phytohemagglutinin (2% v/v, Gibco, cat: 10576015) in supplemented (15% FBS, 100 U/mL penicillin, 100 mg/mL streptomycin) RPMI media. After 48 hours of culture, colcemid (0.1 µg/mL, Gibco, cat: 15212012) was added to stop division (for an additional 2 hours). The cells were treated with a hypotonic KCl solution (0.075 M) for swelling, and chromosomes were fixed with methanol:acetic acid (3:1) and stained with Giemsa. One hundred metaphases on four coded slides were analyzed at 1000x magnification.

Chromosome aberrations were analyzed in metaphases with 44–47 chromosomes, according to international standards ^20^. Chromatid breaks and chromosome fragments, exchanges, translocations, rings and dicentrics were evaluated. Their summed value is the total aberration value. On the basis of decades of practice, 5 aberrations/100 cells are considered the normal range of total aberrations in our laboratory ^21^. The aberrant cell number is the number of cells with any aberration/100 cells, which indicates the frequency of damaged cells.

Chromosomal aberrations were measured before therapy; immediately after treatment; and 3, 6, 9, 12, 24, 36, 48, and 60 months after radiotherapy.

### Statistical analysis

Chi-squared test for categorical and Kruskal-Wallis test for continuous variables were performed to compare the baseline characteristics of patients in different treatment arms. PSA was compared between treatment groups with applied Kruskal-Wallis tests (with Benjamini Hochberg correction). We applied Kruskal-Wallis tests (with Benjamini Hochberg correction) also to analyze differences between follow-up points of chromosome aberrations, IPSS, QoL and baseline values or control group. One sample Wilcoxon test (two-tailed with Bonferroni correction) was used to test if there was a difference between the total aberration values and the normal range of our laboratory. We used multivariate regression analysis to investigate the effects of baseline characteristics on chromosome aberrations. Multivariate linear regression analysis was applied to investigate whether different chromosome aberrations directly after radiotherapy are independent predictors of different side effects in addition to V_100%_ (the volume irradiated with 100% of the prescribed dose) and BED ^22^. All tests and models were considered significant when p < 0.05. For data representation and analysis, StatSoft STATISTICA 12, IBM SPSS Statistics 25, GraphPad Prism (San Diego, CA, USA) and Origin Pro 8.5 were used.

## Results

We followed 25 patients in the conventional LINAC treatment arm (LINAC), 51 patients in the CyberKnife, 68 patients in the low-dose-rate (seed) brachytherapy group and 52 patients in the high-dose-rate brachytherapy treatment (HDR) group. The median follow-up was 60 months in each group.

### Changes in PSA values after radiotherapy

In each treatment group, the average PSA concentration in patients’ blood fell below the upper value of the reference range of our institute (3 ng/mL). However, for HDR-treated patients, the average value was significantly higher than the values after conv. LINAC therapy at all time points for five years (Supplementary figure 1A). Although our sample size was not sufficient to draw conclusions regarding survival, we observed lower biochemical relapse free survival in this group (p=0.004, Supplementary figure 1B).

### Time-dependent chromosome aberrations

An increase in chromosome aberrations due to irradiation was observed in all groups, but we detected 1.7–3.2 times more total aberrations in the groups receiving teletherapy (17.2 ± 2.4/100 cells LINAC, 11.5 ± 1.2/100 cells CyberKnife three months after therapy) than in the brachytherapy groups (6.6 ± 0.5/100 cells LDR, 5.4 ± 0.7/100 cells HDR BT) (Figure 1A). HDR therapy caused the fewest aberrations. After an initial spike, aberrations started to be eliminated, but 60 months later, total aberrations still differed significantly from baseline values (p=0.019 for LDR BT; p=0.046 for LINAC). In our laboratory, 5 total aberrations/100 cells constitute the upper limit for a negative chromosomal aberration test when screening staff at risk of radiation exposure ^21^. The mean total number of chromosomal aberrations in the groups decreased under this limit 24 months after CyberKnife therapy. Directly after LINAC therapy, 3, 6, 12, 24 months later total aberrations were above this limit (one-sample test). Twelve months after LDR BT, the total number of aberrations was significantly greater than 5/100 cells. The median total number of aberrations after HDR was never greater than this threshold, not even immediately after implantation. However, these values were significantly greater than the average total aberration values of the HDR patients before therapy. We also plotted the average total aberrations of our twenty-two age-matched control subjects on the plots (measured once). The total aberration value of all radiotherapy modalities differed significantly from the value of our controls even at baseline except before and directly after LDR brachytherapy.

**Figure 1.**
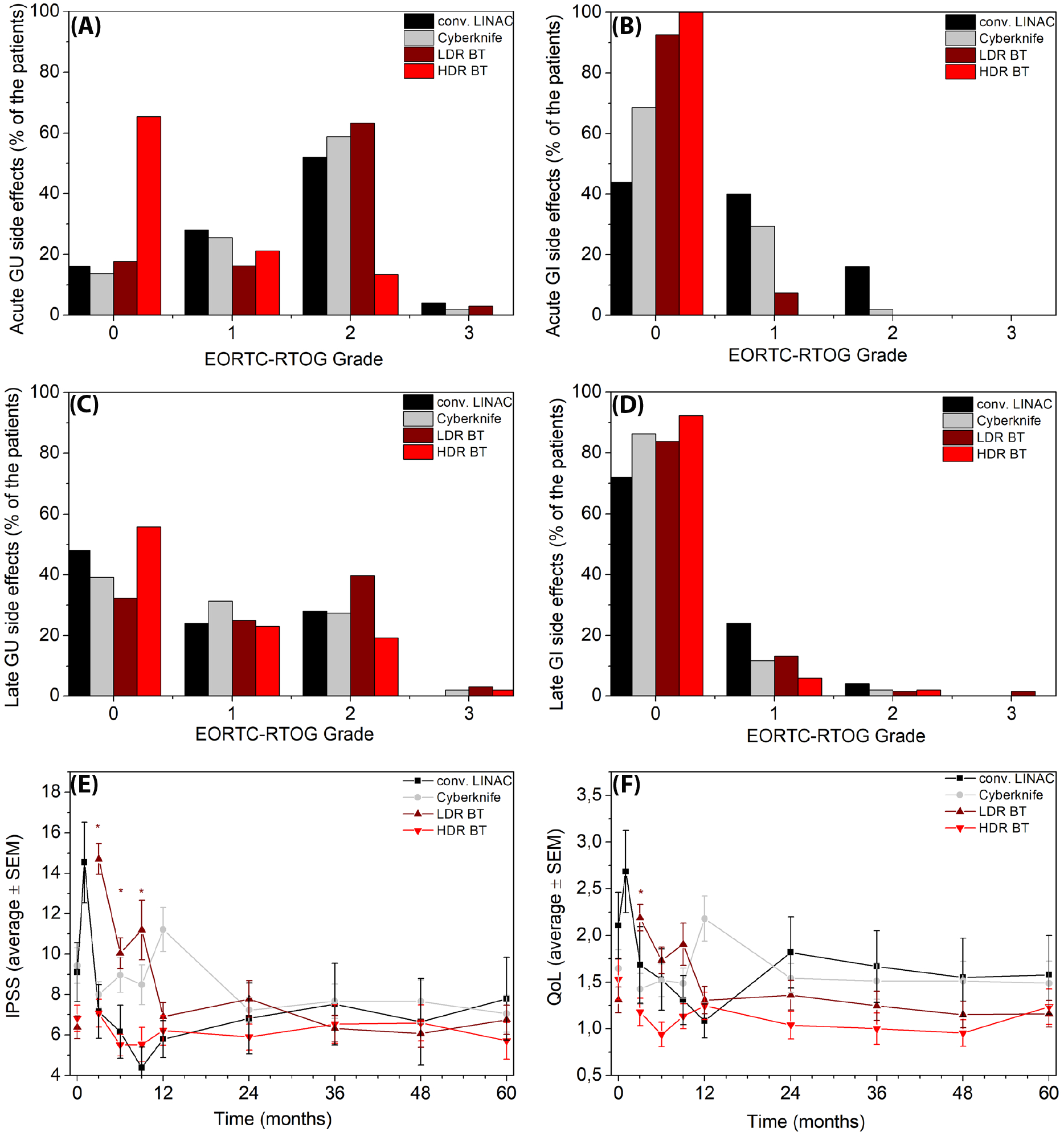
Chromosomal aberrations after conventional LINAC (black) and CyberKnife (gray) therapy, LDR BT (burgundy), HDR BT (red) and age-matched controls (cyan). The time dependence of A) total aberration, B) aberrant cell, C) dicentrics + rings, and D) chromosome fragment frequencies are shown. The significant differences compared with the baseline value of each modality are indicated with asterisks. The error bars represent the SEM.

The time dependence of the aberrant cell frequencies was similar to that of the total aberration values (Figure 1B). Three months after LINAC and CyberKnife therapy, approximately 12.7 ± 1.6% and 8.7 ± 0.8% of the lymphocytes suffered chromosome damage, respectively. Brachytherapy techniques affected fewer lymphocytes (5.3 ± 0.4% for LDR and 4.6 ± 0.6% for HDR therapy at 3 months).

Dicentric and ring aberrations (dicentrics + rings) are radiation-specific chromosomal aberrations that usually do not arise due to environmental conditions ^23^. These values differed significantly from the baseline values at every time point for five years after CyberKnife therapy, four years after conventional LINAC therapy, from three months to five years after LDR-BT, and twelve months after HDR-BT. The dicentrics + rings value was 2.5–7.1-fold greater three months after teletherapy (7.8 ± 1.1 for conventional LINAC, 5.2 ± 0.6 for CyberKnife) than in the brachytherapy arms (2.1 ± 0.2 for LDR BT, 1.1 ± 0.2 for HDR BT). There was no difference between the baseline value of the patients (in any of the radiotherapies) and the dicentrics + rings value of the controls (Figure 1C).

The chromosome break values show a similar trajectory to the values of total aberrations. These aberration values are lower after therapy but persist for years (Figure 1D).

### The role of baseline and treatment characteristics

We performed multivariate regression analysis to test the effects of baseline and treatment characteristics on chromosome aberrations. We decided to include all the characteristics but omit risk, as it is collinear with T status, iPSA and Gleason score, by definition. There was no significant model in the analysis of chromosome aberrations at 3 months (Supplementary Table 2).

### Side effects after different types of prostate cancer radiotherapy

EORTC-RTOG Grade 2 or higher early urogenital side effects occurred in 56.0%, 60.8%, 66.2%, and 13.5% of the patients four years after LINAC, CyberKnife, LDR and HDR therapy, respectively. Of these, only 4.0%, 2.0% and 2.9% were Grade 3 toxicities (LINAC, CyberKnife, and LDR groups, respectively) (Figure 2a). There were no Grade 3 acute gastrointestinal side effects. Furthermore, the frequency of Grade 2 early gastrointestinal effects was 16.0% after LINAC and 2.0% after CyberKnife therapy, and there were no Grade 2 side effects after HDR or LDR BT (Figure 2b).

**Figure 2.**
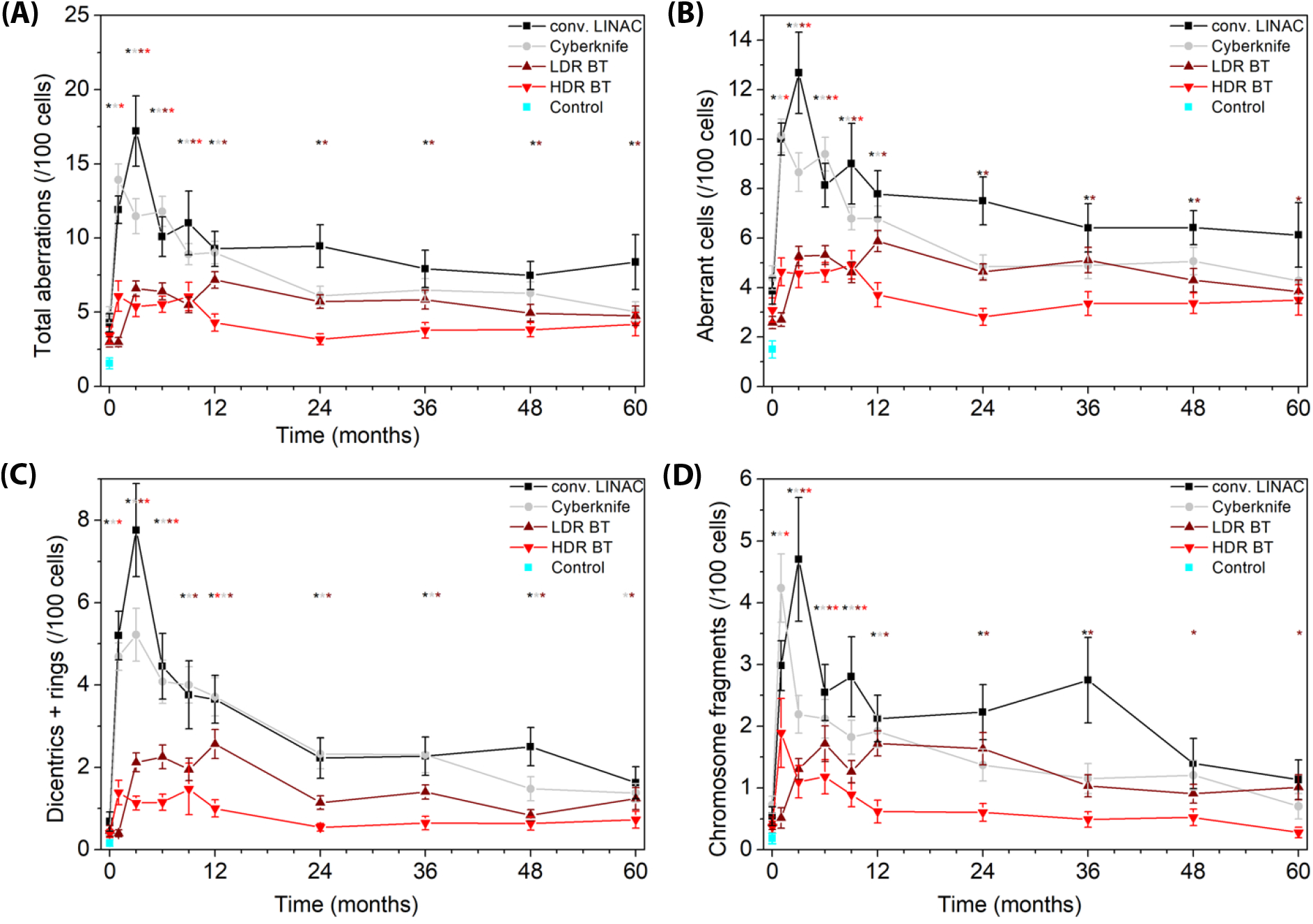
Side effects of conventional LINAC (black) and CyberKnife (gray) therapy, LDR BT (burgundy), HDR BT (red) and age-matched controls (cyan). A) Acute GU; B) acute GI; C) late GU; D) late GI-grade side effects; E) International Prostate Syndrome Score; F) Quality of Life questionnaire scores are shown. The significant differences compared with the baseline value of each modality are indicated by asterisks. The error bars represent the SEM.

With respect to late urogenital side effects, 28.0% were Grade ≥2 after LINAC, 29.4% after CyberKnife, 42.6% after LDR and 21.1% after HDR radiotherapy. The frequencies of Grade 3 toxicity were 2.0%, 2.9% and 1.9% after CyberKnife, LDR and HDR irradiation, respectively (Figure 2c). There was only one (1.5%) Grade 3 late gastrointestinal side effect among the LDR-BT patients, and the percentage of Grade 2 late GI side effects was 4.0% after conv. LINAC, 2.7% after CyberKnife therapy, 1.5% after LDR-BT and 1.9% after HDR-BT (Figure 2d).

We applied patient-reported questionnaires (PROMs) to assess side effects further. The International Prostate Syndrome Score questionnaire assessing urinary toxicities (score range: 0–35, where a higher score indicates worse side effects) was completed by patients at each follow-up visit (Figure 2E). However, it was not appropriate to administer the questionnaires directly after brachytherapy and CyberKnife treatments because the questions asked about the symptoms of the past month. We measured an increase three, six, nine months after LDR therapy (6.4 ± 0.6 to 14.7 ± 0.8 at three months), but no significant difference was found after HDR (6.8 ± 0.6 to 7.1 ± 0.7 at three months), Cyberkinife (9.4 ± 0.9 to 8.0 ± 0.6 at three months) and conventional LINAC therapy (9.1 ± 1.5 to 14.5 ± 2.0 directly after therapy) compared with baseline.

We tested patients’ subjective quality of life (QoL) (score range: 0–6, where a higher score indicates worse side effects), and the scores rose three months after seed therapy (1.3 ± 0.1 to 2.2 ± 0.1), similar to the IPSS results (Figure 2F). There was no increase in HDR, conv. LINAC or CyberKnife scores.

### Connections between chromosomal aberrations and side effects

We performed a multivariate linear regression analysis to study the connection between chromosomal aberrations and side effects to investigate the predictive power. We included biological effective dose and V_100%_ (volume that received 100% of the prescribed dose) in the models in addition to chromosomal aberrations. We chose these variables to balance physical differences in therapy and test if chromosome aberrations can represent individual radiosensitivity. Significant models were identified (Table 2): Total aberration value (p=0.027) and aberrant cell frequency (p=0.015) were independent significant predictors of cumulative genitourinary side effects in addition to V_100%_ and BED.

**Table 2:** Models predicting side effects.

|  | type of side effect | p value (whole model) | Cohen's f<sup>2</sup> (effect size) | p value for the aberration |
| --- | --- | --- | --- | --- |
| Chromosome breaks | GU | 0.039 | 0.045 | 0.497 |
| Total aberrations | GU | 0.005 | 0.070 | 0.027 |
| Aberrant cells | GU | 0.003 | 0.075 | 0.015 |
| Dicentrics +rings | GU | 0.044 | 0.044 | 0.665 |
| Chromosome breaks | GI | 0.647 | 0.099 | 0.942 |
| Total aberrations | GI | 0.623 | 0.099 | 0.735 |
| Aberrant cells | GI | 0.537 | 0.011 | 0.469 |
| Dicentrics +rings | GI | 0.647 | 0.099 | 0.943 |

Multivariate linear regression models with the variables as follows: chromosome aberrations directly after radiotherapy, BED (biological effective dose) and $V_{100\%}$ (the volume irradiated with 100% of the prescribed dose)

## Discussion

To our knowledge, there is no complex analysis of lymphocyte damage after different radiotherapy modalities. We found that the damage to the lymphocytes did not clear from the blood over a period from nine months to five years (depending on the type of therapy). The frequency of damaged cells in the case of teletherapies was greater than 10% at 3 months and approximately 6% after 3–5 years. The difference among radiotherapy techniques is interesting because we could measure the summarized effect of dose rate, overall treatment time, energy, fractionation and other parameters, which differ between the techniques and difficult to calculate.

On the other hand, the clinical significance of these findings needs further investigation. The chromosomal aberration method is performed on lymphocytes, which are induced to divide by phytohemagglutinin M (PHA). According to the literature, mostly T lymphocytes are sensitive to PHA ^24^. One implication of our study is the impact of radiotherapy on the efficacy of immunotherapy, as T-cell DNA damage was shown to have the ability to cause senescence ^4^, which can cause decreased immune surveillance ^6^, promote immunosuppressive phenotype in the tumor microenvironment ^5^ and defects in immunotherapy efficiency ^7-9^. Furthermore, telomere damage can cause T-cell exhaustion as well ^25^. Studying lymphopenia was not among our goals, but we examined the available blood test data and found only 19 lymphocyte counts under 1000/µl in 690 measurements (2.7%). Others also described rare decreases in patients without pelvic irradiation ^26,27^. One can speculate that immunological effects of radiotherapies, which do not affect bone marrow or pelvic lymph nodes, might not be due to lymphocyte count decrease. However, further studies are needed to investigate which immune cell subpopulations are affected. Our work suggests that the use of brachytherapy may hinder immunotherapy less than the use of teletherapies.

The lifetime of lymphocytes remains a subject of debate in the literature. For instance, one study estimated the lifespan of lymphocytes to be 530 ± 64 days in a cohort of 25 women, whereas an upper limit of 29 months has also been proposed ^28^. In another study, chromosome aberrations— which are typically lost during cell division—were still detectable in CD45RA T cells ten years after radiotherapy ^29^. Additionally, studies in mice have demonstrated that naïve T cells can exhibit exceptionally long lifespans ^30^. These findings suggest the existence of distinct lymphocyte subpopulations with differing longevity. In the present study, the inducing reagent phytohemagglutinin M selectively stimulates T lymphocytes, and therefore the measurements primarily reflect this cell population. Moreover, lymphocytes can migrate into lymph nodes, where they may be retained for extended periods ^31,32^. Subsequent re-entry of these cells into the peripheral blood might contribute to an increase in detectable chromosome aberrations at later follow-up time points.

Few publications have applied the chromosomal aberration technique, and even fewer have applied long-term follow-up of radiotherapy patients because the method is highly labor-intensive. However, there are already published studies with similar conclusions, but mostly with shorter follow-up periods. Hille et al. reported significantly more chromosomal aberrations one year after prostate LINAC radiotherapy than before therapy ^13^. They also measured total aberration values after LINAC therapy, which were comparable with our results. Hartel et al. found that one year after radiotherapy for prostate cancer, not just translocations, which are considered long persisting aberrations but also non-transmissible aberrations detected by the FISH technique (dicentrics and non-transmissible exchanges) were more frequent than those at baseline ^33^. Previously, in highly exposed plutonium workers, complex chromosomal aberrations persisted for decades ^34^. Furthermore, noncancer patients irradiated with previous radiotherapy techniques and analyzed via a highly similar chromosomal aberration method presented higher aberration values than did controls approximately 20–40 years after irradiation ^35,36^. These studies included a greater number of patients (max. 143), but in these studies bone marrow could have been affected.

Lymphocytes were used as surrogate normal tissue markers and have been proposed as predictive radiosensitivity markers ^10-12,37^. However, the response to radiotherapy is also dependent on the prescribed dose. We found that HDR therapy, which caused the fewest chromosomal aberrations, also had fewer side effects, but it also caused a slower decrease in PSA and shorter biochemical relapse-free survival (even, if the patient number is insufficient for firm conclusion). In this study, 19 or 21 Gy was given in one fraction as monotherapy, but since the start of our study, 2 × 13.5 Gy has been found to be superior and to provide excellent tumor control ^38^. This is clearly a limitation of our study. On the other hand, randomized studies comparing 21 Gy HDR brachytherapy with other fractionation schedules or LDR brachytherapy have not been performed. LDR brachytherapy also caused fewer chromosomal aberrations than did teletherapies but is known to lead to similar biochemical relapse-free survival. Our analysis was also limited by the fact, that we analyzed smaller number of conventional LINAC patients, however, patient preference shifted toward faster therapies. There was an initial imbalance in IPSS values, the conv. LINAC and CyberKnife group had more initial mean IPSS than brachytherapy groups but the initial imbalance was less than the difference after the therapies.

We also investigated the connection between chromosomal aberrations and side effects to shed light on their predictive value. Lymphocyte chromosome aberrations do not cause toxicities but serve as surrogate predictors. We corrected the analysis with the biologically effective dose and irradiated volume (V_100%_) and found that total aberration and aberrant cell values are independent predictors of cumulative genitourinary toxicities. According to our previous publication, there is a direct correlation between in vivo aberrations measured after radiotherapy and aberrations in the in vitro irradiated blood of these patients (taken before therapy) ^39^. These results may suggest that the chromosomal aberration assay may be suitable for selecting radiosensitive patients. On the other hand, the method is not a high-throughput technique, and other predictive tests may be more likely to be introduced into clinical use ^40^.

We analyzed nearly 196,000 cells one by one under a microscope (196 patients, 10 samples each, at least 100 cells/sample). There are only a few chromosomal aberration studies (with long follow-up periods) in the literature because the method is highly labor intensive. We also described the effect of CyberKnife therapy, which has emerged as a new therapeutic modality, and there is a need to identify its niche in prostate therapy ^41^.

## Conclusions

We found that, compared with brachytherapy, teletherapy increased the chromosome aberrations of the lymphocytes by 1.7–3.2 times after prostate radiotherapy. The number of affected lymphocytes can reach 13% in the first 3 months. In our study, radiation-induced chromosomal aberrations in lymphocytes did not disappear (return to baseline values) even 5 years after therapy. Insight into the subtypes of damaged lymphocytes is needed to assess their implications.

## Supporting information

Supplementary Figure 1 subtext

## List of abbrevations

BT: brachytherapy
CTV: clinical target volume
EORTC-RTOG: European Organization for Research and Treatment of Cancer-Radiation Therapy Oncology Group
FBS: fetal bovine serum
GI: gastrointestinal
GU: genitourinary
GS: gleason score
HDR: high dose rate
HT: hormone therapy
IPSS: International prostate symptom score
LDR: low dose rate
LINAC: linear accelerator
PROM: patient-reported outcome measure
PTV: planning target volume
SIB: simultaneous integrated boost
QoL: quality of life
V_100%_: volume, which received the 100% of the prescribed dose

## Conflict of interest

There are no financial or nonfinancial competing interests.

## Data availability statement

The datasets used within the current study will be shared upon reasonable request.

## Ethics statement

We performed the study according to the Helsinki guidelines. The ethical permit number is 16738-2/2015/EKU, approved by ETT TUKEB on 03.24.2015. Both the patients and volunteers signed informed consents.

## Funding

This work was supported by the National Research, Development and Innovation Fund of the Ministry of Culture and Innovation under the National Laboratories Program (National Tumor Biology Laboratory (2022-2.1.1-NL-2022-00010)); the Hungarian Thematic Excellence Program (TKP2021-EGA-44); and the Investment in the Future (Development of Innovative Cancer Diagnostic and Therapeutic Procedures at the National Institute of Oncology (2020-1.1.6-JÖVŐ-2021-00008)) Grant Agreements with the National Research, Development and Innovation Office.

## Acknowledgments

We greatly acknowledge the work of Nagyesda Vass, Krisztina Kiss and Martin Fodor.

## Notes

### Competing Interest Statement

The authors have declared no competing interest.

### Summary of Updates

We update five year data, in all sections

