## Supplementary figures and images for "Radiotherapy Technique Determines the Magnitude and Persistence of Lymphocyte DNA damage"

### Supplementary Figure 1

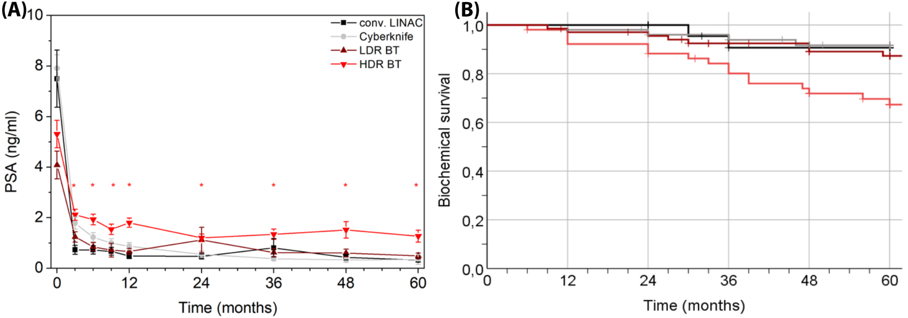
