## Supplementary Figure 1 subtext for "Radiotherapy Technique Determines the Magnitude and Persistence of Lymphocyte DNA damage"

Supplementary figure 1: PSA decrease (A) and biochemical relapse free survival (B) as a result of conventional LINAC (black) and CyberKnife (gray) therapy, LDR BT (burgundy) and HDR BT (red) (A). The significant differences compared with the PSA values after conv. LINAC therapy are indicated with asterisks. The error bars represent the SEM.
